# Previously unrecorded distribution of marine sediments derived yeast isolates revealed by DNA barcoding

**DOI:** 10.1101/2020.08.29.273490

**Authors:** Chinnamani PrasannaKumar, Shanmugam Velmurugan, Kumaran Subramanian, S. R. Pugazhvendan, D. Senthil Nagaraj, K. Feroz Khan, Balamurugan Sadiappan, Seerangan Manokaran, Kaveripakam Raman Hemalatha, Wilson Aruni, Bhagavathi Sundaram Sivamaruthi, Chaiyavat Chaiyasut

**Affiliations:** Biological Oceanography Division, CSIR-National Institute of Oceanography, Dona Paula, Panaji, Goa-403004, India; Centre of Advance studies in Marine Biology, Annamalai University, Parangipettai, Tamil Nadu-608502, India; Madawalabu University, Bale, Robe, Ethiopia; Centre for Drug Discovery and Development, Sathyabama Institute of Science and Technology, Tamil Nadu-600119, India; Department of Zoology, Arignar Anna Government Arts College, Cheyyar, Tamil Nadu-604407, India; Research Department of Microbiology, Sadakathullah Appa College, Rahmath Nagar, Tirunelveli Tamil Nadu -627011, India; Center for Environment & Water, King Fahd University of Petroleum and Minerals, Dhahran-31261, Saudi Arabia; Department of Microbiology, Annamalai University, Annamalai Nagar, Chidambaram, Tamil Nadu-608002, India; Department of Biotechnology, School of Bio and Chemical Engineering, Sathyabama Institute of Science and Technology, Chennai, Tamil Nadu-600119, India; US Department of Veteran affairs, Loma Linda, CA, USA; Innovation Center for Holistic Health, Nutraceuticals, and Cosmeceuticals, Faculty of Pharmacy, Chiang Mai University, Chiang Mai 50200, Thailand

**Keywords:** Marine yeast, DNA barcoding, unrecorded yeasts, yeast pathogens, ITS gene

## Abstract

For the yeast population and diversity, marine habitats are the least explored niches and the lack of validated database is considered to be a drawback for yeast research. The aim of the present study is to create a comprehensive DNA barcode library for marine derived yeast species isolated from organic burial hotspots such as coastal sediment in mangrove and continental shelf ecosystems. As we enriched, isolated and ITS gene sequenced 1017 marine derived yeast isolates belonging to 157 marine species in 55 genera, 28 families, 14 orders, 8 classes of 2 Phyla (*viz*., Ascomycota and Basidiomycota) of which 13 yeast species were first time barcoded. We witnessed yeast species of both terrestrial and marine endemic origin in the barcode datasets. Due to the large volume of sequencing trace files, the variable length of extracted sequences, and the lack of reference sequences in public databases, difficulties were faced in taxonomic sequence validation. The length of the majority of the sequences (99.42%) were more than or equal to 600 base pairs. BLAST analysis revealed that 13 yeast species were barcoded for the first time. The genus, *Candida* was the speciose genera isolated in this study. K2P intra-species distance analysis performed for selective groups yielded an average of 0.33%, well below the previously proposed yeast barcode gap. ITS gene NJ-tree based identification conducted for selective species in Ascomycota and Basidomycota, precisely clustered the same species into one group, indicating the efficacy of ITS gene in yeast species delineation. Besides isolating some of the common marine yeast species such as *Candida oceani, Bandonia marina* and *Yarrowia lipolytica*, we found approximately 60% of the yeast species isolates were previously unrecorded from the marine environment (example; *Cystobasidiopsis lactophilus, Slooffia cresolica, Udeniozyma ferulica, Colacogloea falcatus* and *Pichia guilliermondii*), of which 16.5% were recognised as potential human pathogens (example; *Candida orthopsilosis*, *C. rugosa, Debaryomyces fabryi* and *Yamadazyma triangularis*). Apart from releasing the barcode data in GenBank, provisions were made to access the entire dataset along with meta-data in the Barcode of life database (http://dx.doi.org/10.5883/DS-MYIC). This research constitutes the largest dataset to date for collecting marine yeast isolates and their barcodes. As meta- and environmental barcoding analysis were expanding its scope including environmental assessment and monitoring, the datasets such as ours will be more useful.

## 1. Introduction

In the tropical and subtropical geographies, mangroves and its associated mycobiota are vital components of the ecologically and economically significant food web (Lee et al., 2014). So far 69-77 mangrove species (including their associates and hybrids) have been documented (Alappatt, 2008; Statzell-Tallman et al., 2008). The mangroves constitutes 4 Gton of Carbon with global biomass of 8.7 Gton, distributed in 112 countries over 10-24 million hectares (Kathiresan & Bingham, 2001; Giri, 2011). The primary production rate in mangrove ecosystems is equal to the rates in evergreen tropical humid forests (Alongi et al., 2014). On the other hand, in the Bay of Bengal, one of the largest sediment inputs occurs among the world’s ocean, receiving 2000 million tons of sediments annually (Mohanty et al., 2008) along with significant amount of terrestrial organic materials (Hedges and Keil, 1995; de Haas et al., 2002) contributed by various rivers including the Kaveri on the southeast coast of India (Khan et al., 2012). Yeast populations have rarely been explored in these sites besides an abundant occurrence of culturable filamentous fungal isolates in these large carbon burial sites (Das et al., 2009; Velmurugan et al., 2013; Godson et al., 2014).

In comparison to aquatic ecosystems, fungi in terrestrial and fresh water environments have been well studied (Gulis et al., 2009; Raja et al., 2018). Through decomposition plant material and nutrient replenishments (Schmidt & Shearer, 2004; Shearer et al., 2007), fungi plays a significant role in the mangrove ecosystems by providing food for fishes and invertebrates (Hogarth, 1999). The unicellular fungi, yeast (Kutzman and Fell, 2015) is a polyphyletic group (Kutty and Philip, 2008), and when requires seawater for its growth was described as marine yeast (Chi, 2012). Marine yeasts are known for their parasitic, mutualistic or saprophytic relationship with marine animals in addition to their significance in nutrient cycling, and may therefore be associated with various marine invertebrates, including crabs, clams, mussels, prawns, oysters or other substrates (Jones and Pang, 2012; de Araujo et al., 1995; Kosawa da Costa et al., 1991; Pagnocca et al., 1989). In two major phyla viz., Ascomycota and Basidiomycota, the diverse marine yeast is largely classified (Kurtzman 2011; Boekhout et al. 2011). Though basidiomycetous yeast species were known for high salt tolerance (Tekolo et al., 2010), most of marine derived Ascomycetous yeast species are of terrestrial origin with widespread phylogenetic diversity (Fell, 2012).

Since its first isolation (Fischer and Brebeck, 1894), which was recently overcome using molecular taxonomy (Fell, 2012), numerous difficulties have existed in marine yeast nomenclature and taxonomy. DNA barcoding has simplified the recognition of biological species using short DNA fragment sequencing and analysis (Hebert et al., 2003). One of the vision behind DNA barcoding was the simple identification of biological species by nonexperts for the advancement of biological, ecological and medical research. Internal transcribed spacer (ITS) gene has been recognised as DNA barcode which successfully delineates filamentous fungal species (Schoch et al., 2012; Velmurugan et al., 2013; Vu et al., 2019) and even works better for classification of yeast species (Vu et al., 2016), substituting the previously used laborious and frequently inaccurate phenotypic analysis (Kurtzman et al., 2015). Lack of validated data sets for yeast research is considered to be a drawback for its identification (Bidartondo, 2008). The goal of this study is to use DNA barcoding to explore the marine sediments in mangrove swamps along Indian coast and continental shelf off southeast coast of India for yeast diversity. We aim to synthesise and publish, in GenBank and BOLD, a sizable amount of marine derived yeast DNA barcodes. Besides evaluating such database for identification of yeast species, we suspect that even after the production of large yeast barcode dataset (8669 barcodes for 1351 yeast species) (Vu et al., 2016), marine habitats will still be able to incubate some yeast species yet to be barcoded. The purpose of the study was to gather as many marine derived yeast cultures as possible for a comprehensive DNA barcode library synthesis, as studies are uncommon in exploring such a large scale marine environments for yeast isolates.

## 2. Materials and methods

### 2.1. Study area and sample collection

Between Nov, 2008 and Jan 2013, sediment samples were extensively collected from two separate ecosystems, viz., 1) inter-tidal sediments flats under mangrove trees along Indian’s coastline, 2) continental shelf sediments off India’s southeast coast. The setails of sampling sites with geographical descriptions were given in **Table S1**.

Inter-tidal sediments under mangrove trees were collected along the Indian coastline from various mangrove environments ranging from Gujarat’s Kachchh Mangroves in West coast of India to West Bengal’s Sundraban ecosystem in East coast of India. Sampling of 14 mangrove station called M1 through M14 and details of collection date, geographical coordinates, and number of samples per station are provided in table S1. During pre-monsoon seasons, mangrove sediments were collected mainly at low tide. Approximately 50 to 100g of undisturbed sediments were collected (in triplicates) using sterile spatula, transferred to sterile polythene bags (Nasco bags, HiMedia), and transported to laboratory in ice. Samples were processed normally within 48 hours. In mangrove environment, sediments (n=52) were obtained from 13 mangrove species viz., *Avicennia* sp., *A. ilicifolius, A. alba, A. marina, A. officinalis, Bruguiera cylindrical, Ceriops decandra, Excoecaria agallocha, Pongamia pinnata, Rhizophora mucronata, Sonneratia apectala, S. caseolaris* and *Xylocarpus mekongensis* in 14 sampling stations (named as M1 –M14) (table S1). Sediment pore water salinity was measured using hand-held refractometer (BEXCO, USA). Highest of 35ppt salinity was measured in Parangipettai and Pitchavaram mangroves (station, M9 and M10, respectively) whereas station M14 which is upstream mangroves infringe inland of Sundarbans, West Bengal, records the lowest salinity of 8ppt.

During cruise No. 260 (Lyla et al., 2012) of Fishery and Ocean Research Vessel (FORV) Sagar Sampada conducted from 06^th^ to 28^th^ December 2008 using Smith–McIntyre grab which covers an area of 0.2m^2^, sediment samples along the Southeast continental self of India were collected. On the Southeast continental shelf, the sampling station was fixed at the depth gradients of about 30, 50, 100, 150 and 200 meters at each transacts, constituting a total of 7 transects and 32 sampling stations, called from CS1 through CS32 (**table S1**). In-built with FORV-Sagar Sampada, the depths of sediment sampling were measured using multibeam echo sounder (capable of measuring up to 1000m depth). Sterile spatula was used for sediment sub-sampling (trice per station) from the grab. The cruise collected a total of 96 sediment samples. Salinity of continental shelf sediments ranged from 33 to 35ppt. All culture media were prepared using the ambient seawater (AS) obtained from respective collection sites, as it provides closer to natural environmental conditions (Zaky et al., 2014). AS was filtered through 0.22μ cellulose filter membrane (Miilipore) and autoclaved, before being used to prepare culture media.

### 2.2. Enrichment and isolation of yeast-like cells

Two types of media were used for enrichment, as enrichments were known to increase the number of yeast species isolates (Gadanho & Sampaio, 2004). Either one or both of the media (for most samples collected after 2010) was used for enrichment of yeast cells before plate culture. Briefly, after homogenization of the sediment samples in the collection container, yeast cells were enriched by adding one gram (g) of sediments to 100ml of Yeast/Malt extract (YM) broth (3 g malt extract, 3 g yeast extract, 10 g dextrose, and 5 g peptone, in 1L sterile-AS) and/or GPY broth (2% glucose, 1% peptone, 0.5% yeast extract in 100ml sterile-AS) supplemented with an antibiotic cocktails (300 mg L^−1^ penicillin, 300 mg L^−1^ streptomycin, 250 mg L^−1^ sodium propionate, and 0.02% of chloramphenicol) to inhibit bacterial growth. We used both enrichment media for sediments sampled after 2010, to increase the number of yeast species being isolated. At 150 rpm, the enrichment broth with sediment samples in a 250 mL Erlenmeyer flasks was shaken on a rotary shaker, incubated for 2-3 days at 17-20 °C (temperature >20 °C was found to accelerate filamentous fungal growth in plate cultures). Autoclaved AS has been used as control. After incubation from corresponding broth cultures, 100μl to 1000μl (based on the turbidity of the broth) was spread over (in triplicates) YM and/or GYP agar plates (composition as same for broth preparation with addition of 1.5% agar). Culture media was autoclaved twice (at 100°C for 30min) during two consecutive days to reduce mould contaminations (Gadanho and Sampaio, 2005). The remaining broth was conserved at 4°C with the over lay of mineral oil for future use, just in case the incubated plates did not produce any colonies or over production of filamentous fungi. The inoculated plates were incubated for 10-20 days or until colonies appeared and continuously monitored at every 24 hours. In order to promote the full recovery of yeast-like species including slow growing colonies, prolonged incubation period with concurrent removal of fast growing colonies were adopted. Also care was taken to stop the incubation when there was a high probability of mould over growth.

Microscopic analyses of yeast-like colonies were started from the minimum of 5 days of incubation with methylene blue staining (checked for single type forms and to ensure no association of bacterial cells) and purified by streaking onto fresh agar (YM or GYP) plates (to prevent growth over other colonies or to save from rapidly growing moulds). Gradually one representative morphotype of each colony per sample (i.e., when the yeast colonies were <50 numbers or when the mould was entirely absent or scarce in the agar plates) was streaked twice on corresponding agar plates for purification. The representative number of colonies (40-70 colonies) is randomly selected and purified twice in other cases (i.e., when yeast colonies is >50 or dense mould growth in the agar plates).

Enrichments cultures as mentioned above were done for continental shelf sediments on the ship board Microbiology lab, FORV-Sagar Sampada. After 2 sub-culturing of yeastlike cells, the isolates were grown for 2-3days in the shaker (at 120 rpm) 1mL of GPY broth (in duplicates) prepared in 2mL microfuge tubes. The culture in one tube was used for DNA isolation and other cryopreserved (culture increased to 1.5ml volume using 2% GPY broth, 10% glycerine) in the Marine Microbial Culture Facility, Centre of Advanced Study in Marine Biology, Annamalai University. There were a total of 1398 colonies for molecular analysis (916 colonies from mangrove sediments, and 482 colonies from continental shelf sediments). Each purified cultures were numbered under DBMY (DNA Barcoding Marine Yeast) acronym. The key features described in Yarrow (1998) and Kurtzman et al. (2010) were used for the macro- and micro-morphological analysis for identification.

### 2.3. DNA extraction, polymerase chain reaction and DNA sequencing

In order to recover the yeast-like cells grown in the 2mL microfuge, the tubes were centrifuged at 8000 X g for 5 minutes. Following manufacturer’s instructions, DNA was extracted from the pellets using GeneiPure Yeast DNA preparation kit (GeNei) or DNeasy blood and tissue kit (Qiagen). The DNA was eluted in the elution buffer (provided with the kit) and stored at −20°C. The ITS primers; ITS1 (5’-TCCGTAGGTGAACCTGCGG) and ITS4 (5’-TCCTCCGCTTATTGATATGC) (White et al., 1990) was used for amplification. The primers targets the DNA fragments containing, partial 18S ribosomal RNA gene; complete sequences of internal transcribed spacer 1, 5.8S ribosomal RNA gene, internal transcribed spacer 2 and partial sequences of 28S ribosomal RNA gene. PCR was performed on a thermal cycler 130045GB (GeNei) under following conditions: 4 min at 94 °C, followed by 30 cycles of 30 s at 94 °C, 40 s at 48 °C for annealing and 90 s at 72 °C, with a final extension at 72 °C for 7 minutes. PCR amplicon were separated by 1.5% agarose gel electrophoresis. Amplicons were sequenced two ways using commercial sequencing services of Macrogen (South Korea) or Bioserve Biotechnologies Pvt. Ltd. (India).

### 2.4. DNA sequence analysis

Following sequencing, the forward and reverse sequences were assembled using BioEdit ver. 5.0 (Hall, 1999). Sequences were aligned using CLUSTAL X (Larkin et al., 2007) and manually adjusted in MEGA X (Kumar et al., 2018). DNA sequences were compared with GenBank sequences using BLAST algorithms (Altschul et al., 1997) and a cut-off species threshold of 98.41% (Vu et al., 2016) was used for delamination of yeast species. The species that were first time barcoded (i.e., when species threshold is <98%) were confirmed by double checking BLAST search similarity values and by searching for ITS gene sequences of the species in GenBank.

Using the reference sequences extracted from GenBank, the Neighbor-Joining method (Saitou and Nei, 1987) was used for tree based yeast species identification. In the bootstrap test, the percentage of replicate trees in which the associated taxa clustered together (100 replicates) (Felsenstein, 1985) is indicated as circles next to the branches. The tree is drawn to scale, with branch lengths in the same units as those used to infer the phylogenetic tree from evolutionary distances. The evolutionary distances have been computed using the Kimura 2-parameter method (Kimura, 1980) and are in the units of the number of base substitutions per site. All positions containing gaps and missing data were eliminated (complete deletion option). Evolutionary analyses were conducted in MEGA X (Kumar et al., 2018). The NJ trees were manipulated in interactive Tree of Life (iTOL) database (Letunic and Bork, 2019) for better representation.

DNA sequences generated in the present study were release to GenBank and could be accessed through accession numbers: KJ706221-KJ707237. Entire dataset produced in this study could also be accessed through Barcode of life database under the project title “DNA barcoding marine yeast” with a tag, “DBMY” or through a digital object identifier; http://dx.doi.org/10.5883/DS-MYIC.

## 3. Results and discussion

### 3.1. Composition of marine yeast species revealed through BLAST analysis

Out of the 1398 isolates selected for molecular analysis, 1017 (72.74%) were successfully sequenced. The BLAST analysis shows that ITS gene sequences belonged to 157 yeast species in 55 genera, 28 families, 14 orders, 8 classes (viz., Agaricostilbomycetess, Cystobasidiomycetes, Dothideomycetes, Exobasidiomycetes, Microbotryomycetes, Saccharomycetes, Tremellomycetes, Ustilaginomycetes) in 2 Phyla (*viz*., Ascomycota and Basidiomycota). Two species (*viz*., *Curvibasidium cygneicollum* (J.P. Sampaio, 2004), and *Hasegawazyma lactosa* (T. Hasegawa, 1959)) could not be classified into any defined family, and were therefore placed under *incertae sedis*. Of the total number of sequences (n=1017) generated, 52.01% (n=529) of the sequences were the representatives of Ascomycota and the remaining 47.99% (n=488) of the sequences were the representatives of Basidiomycota. List of recovered species and their respective sampling stations were given in **Table 1**.

**Table 1:**
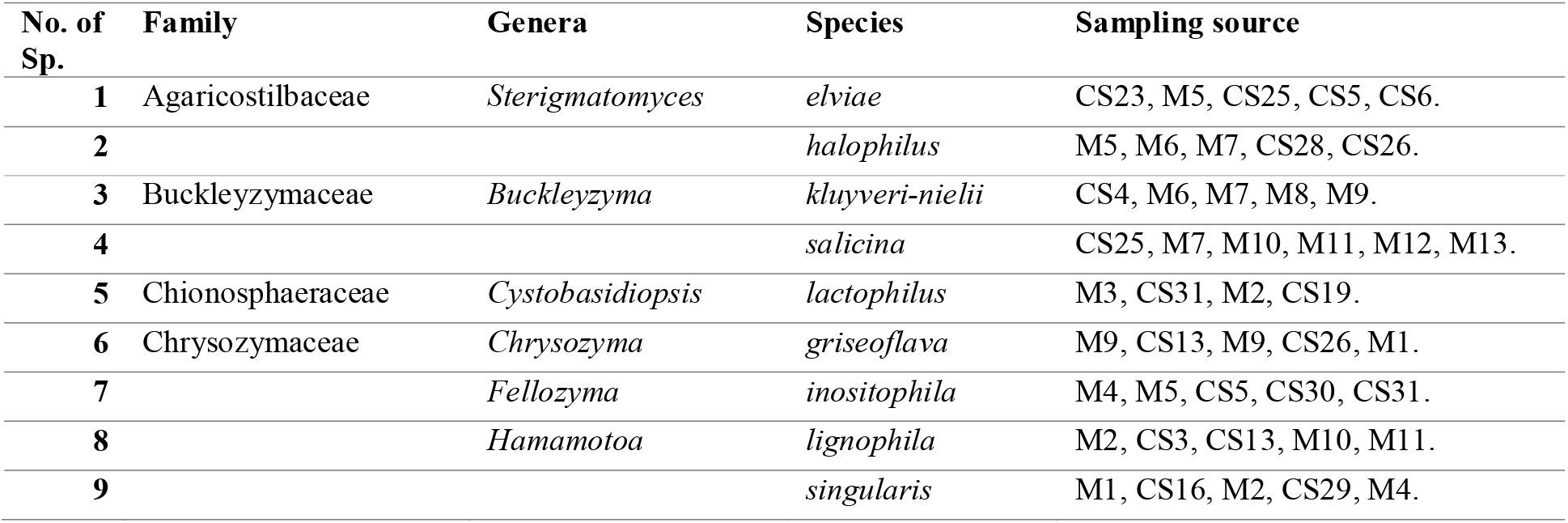
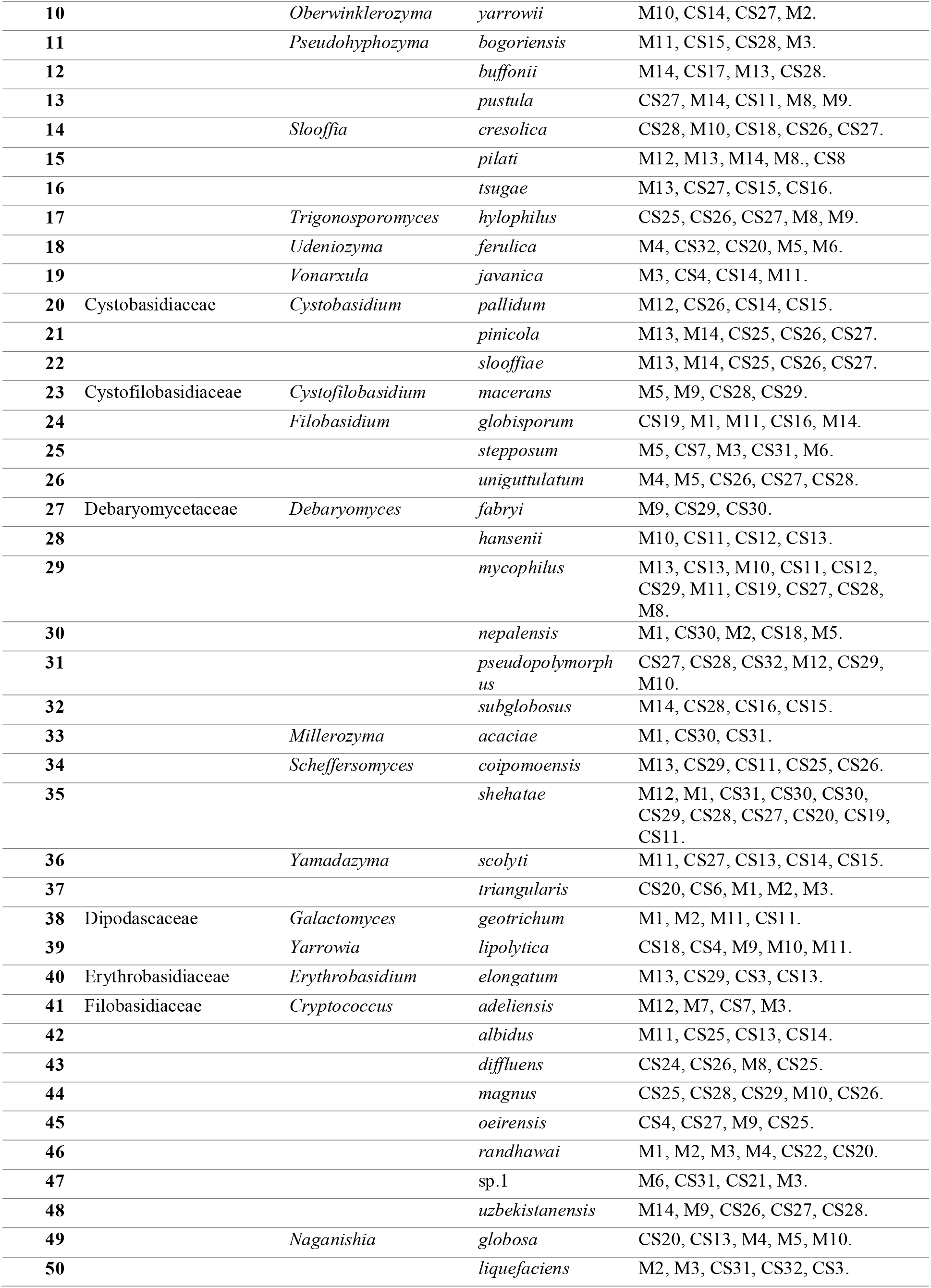
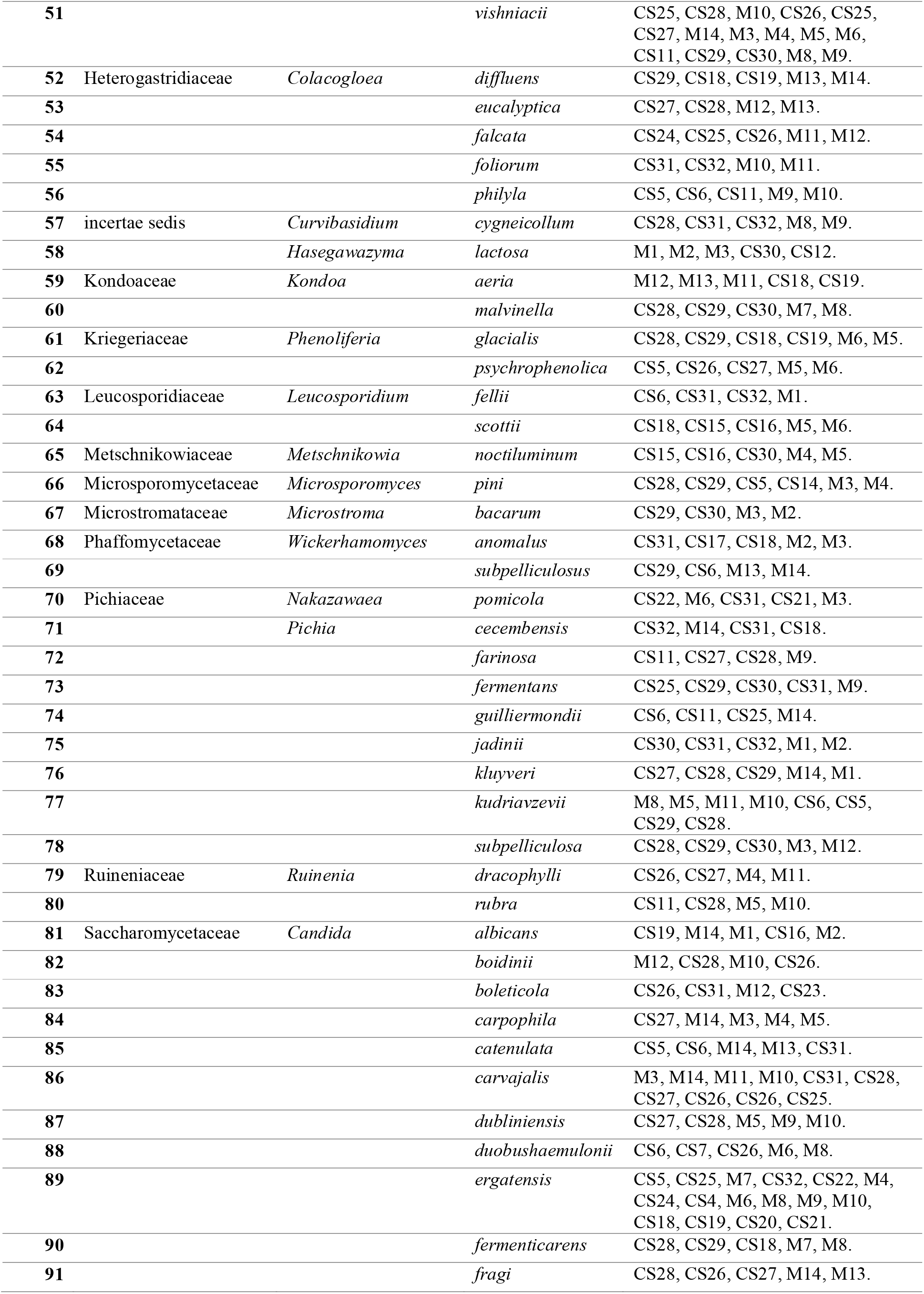
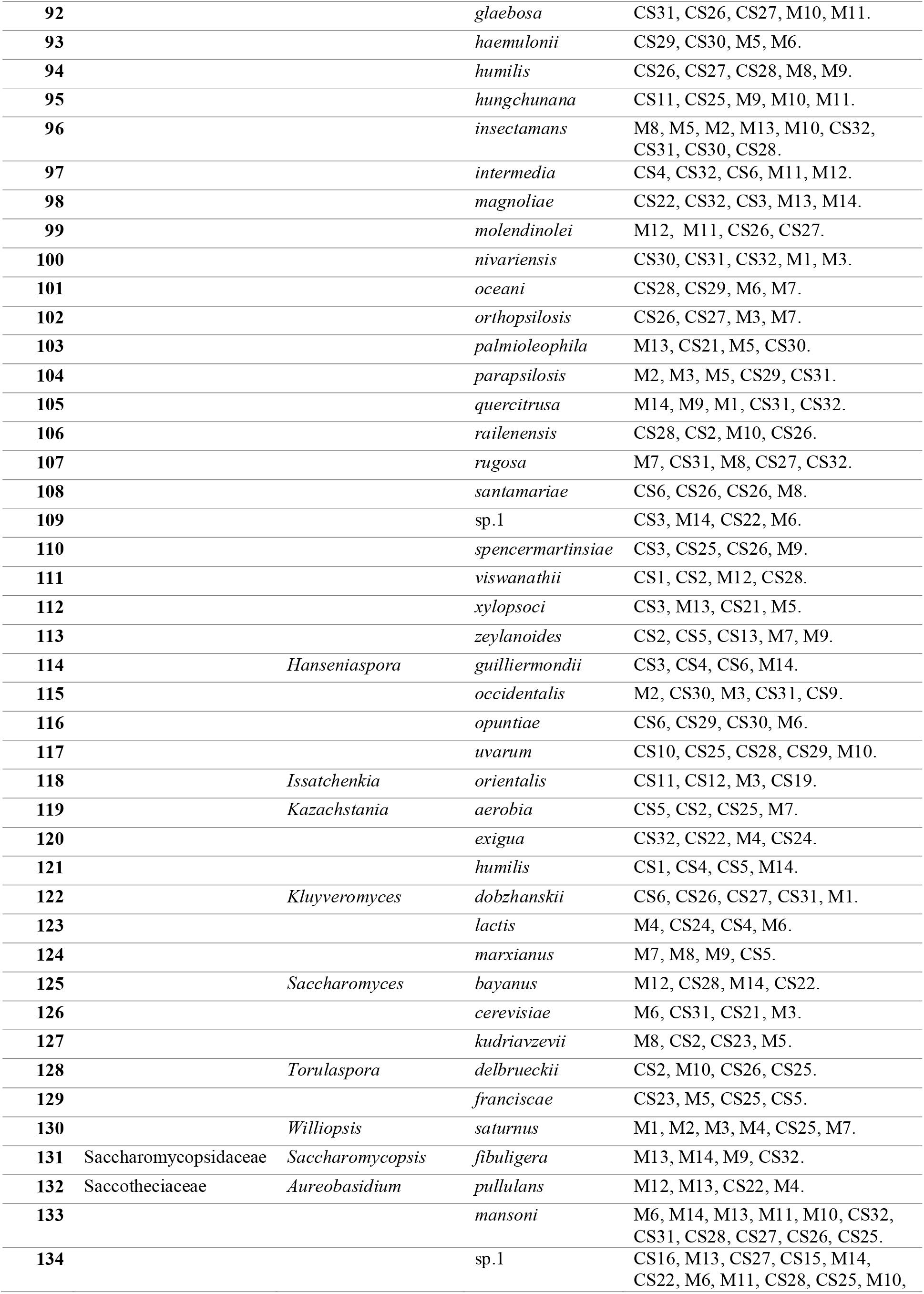
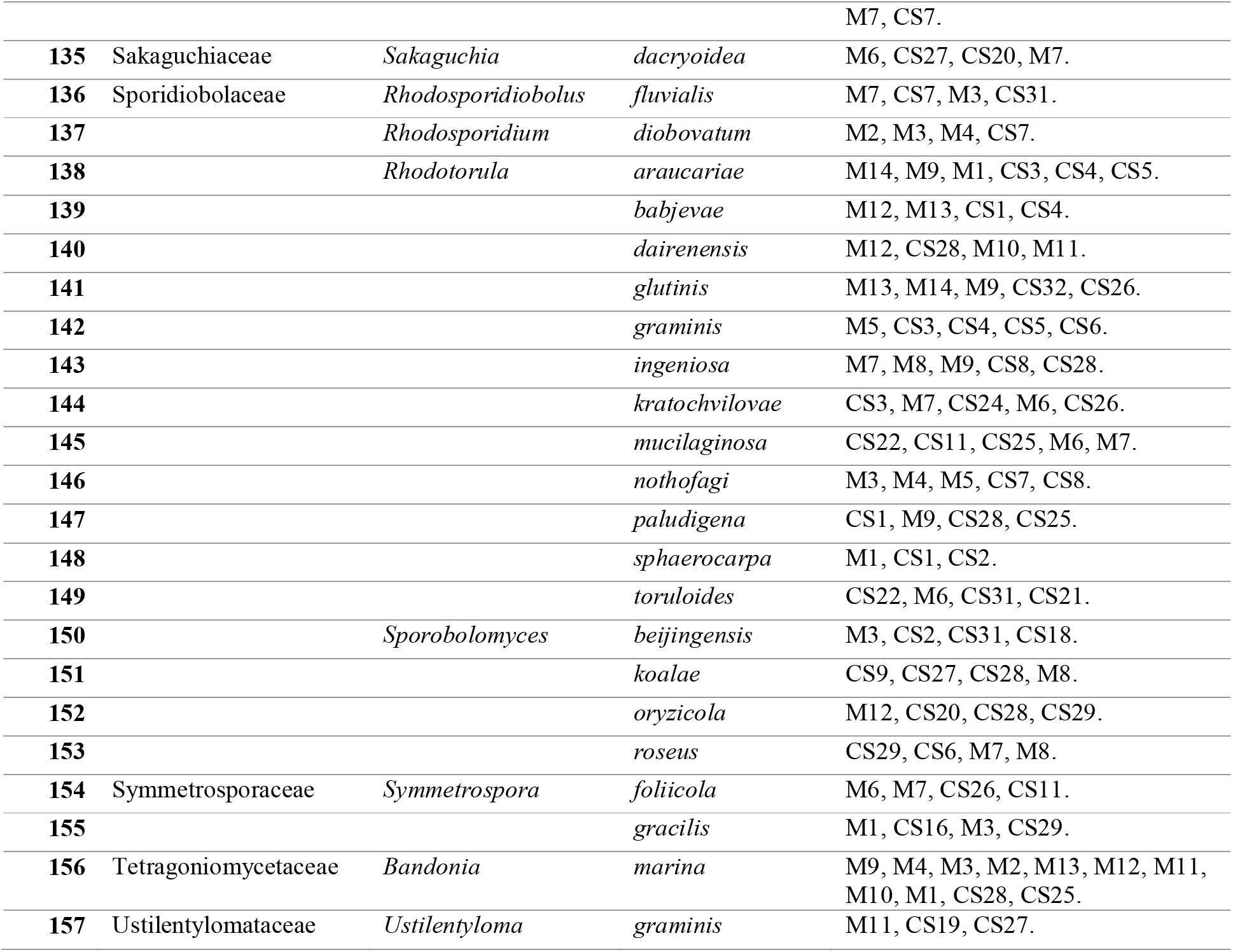
List of marine derived yeast species and its respective sampling station. Colonies that could not be identified to species level was indicated by “sp.”

Minimum of 6 isolates was sequenced from each yeast species isolated in this study. The isolates of *Scheffersomyces ergatensis* (Santa María, 1971), *Naganishia vishniacii* (Vishniac & Hempfling, 1979) and *Aureobasidium* sp. (Viala & G. Boyer, 1891) provided a maximum of 19 sequences each. Each of the yeast species; *viz*., *Aureobasidium mansonii* (Cooke W.B., 1962), *Bandonia marina* (Uden & Zobell, 1962), *Candida carvajalis* (James S. A., 2009), *Candida insectamans* (D.B. Scott, Van der Walt & Klift, 1972), *Debaryomyces mycophilus* (Thanh, Van Dyk & M.J. Wingf., 2002), *Scheffersomyces shehatae* (H.R. Buckley & Uden, 1964) was sequenced for 13 times and the *Pichia kudriavzevii* (Boidin, Pignal & Besson, 1965) was sequenced for 12 times. *Candida* spp. was the most speciose genera (n=33 species) followed by *Rhodotorula* spp. (n=13 species). Largely, we did not find clear pattern of differentiation in yeast species isolated from mangrove and continental shelf sediments.

### 3.2. Character based identification and BLAST analysis

The length of the ITS sequence recovered varied from 542bps to 891bps (**Fig. 1**). The majority of the sequences (62.24%; n=633 sequences belonging to 81 species) were between 600 and 649 bps. Minimum length of 552 - 599 bps was only recovered only for 6 sequences (0.58%). All remaining recovered sequences (99.42%) were larger than or equivalent to 600bp length. The list of 1017 barcodes with their respective species match was given in **Table S2** containing details of the GenBank reference sequence (with percentage of similarity, its accession numbers and its taxonomy).

**Fig. 1:**
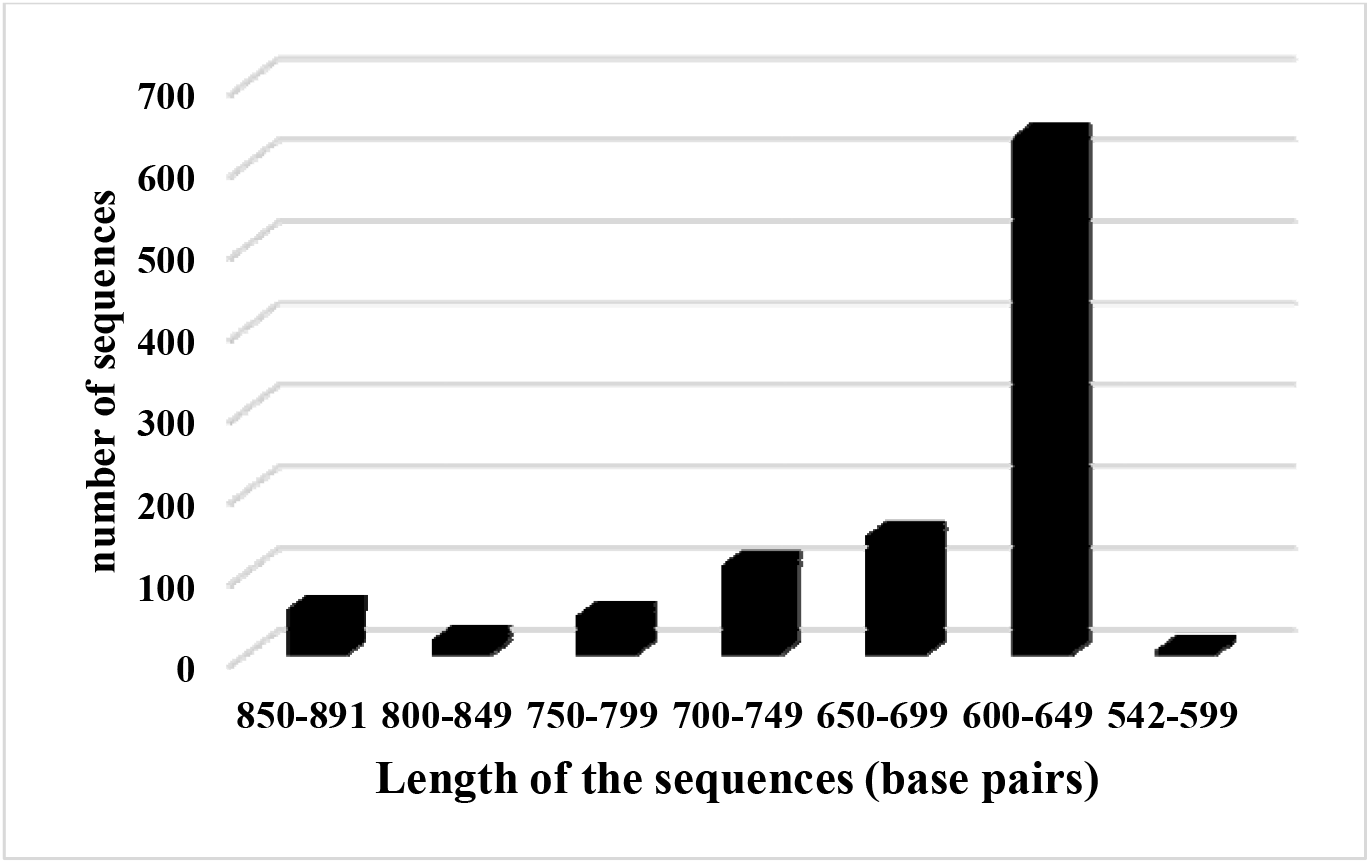
ITS gene sequence length (bps) variations distributed among 1017 sequences. More than 60% of the sequences were in between 600 and 649 bps lengths.

BLAST analysis revealed that 13 yeast species were barcoded for the first time (**Table 2**) and individual search of the ITS gene of those species in GenBank did not yield any results, re-confirming that those species were first time barcoded. Despite *Candida* spp. was speciose genera reported and DNA barcoding, the ITS gene sequences of *Candida carvajalis* (n=15), C. *duobushaemulonii* (n=6) and *C*. *haemulonis* (n=6) was barcoded for the first time, as they were absent in the reference database until now. Even though previous extensive study, barcoded 1351 yeast species producing 8669 barcodes (Vu et al., 2016), the above mentioned 13 *Candida* spp. species was not included in their collection, as the previous study did not explore marine environments.

**Table 2:**
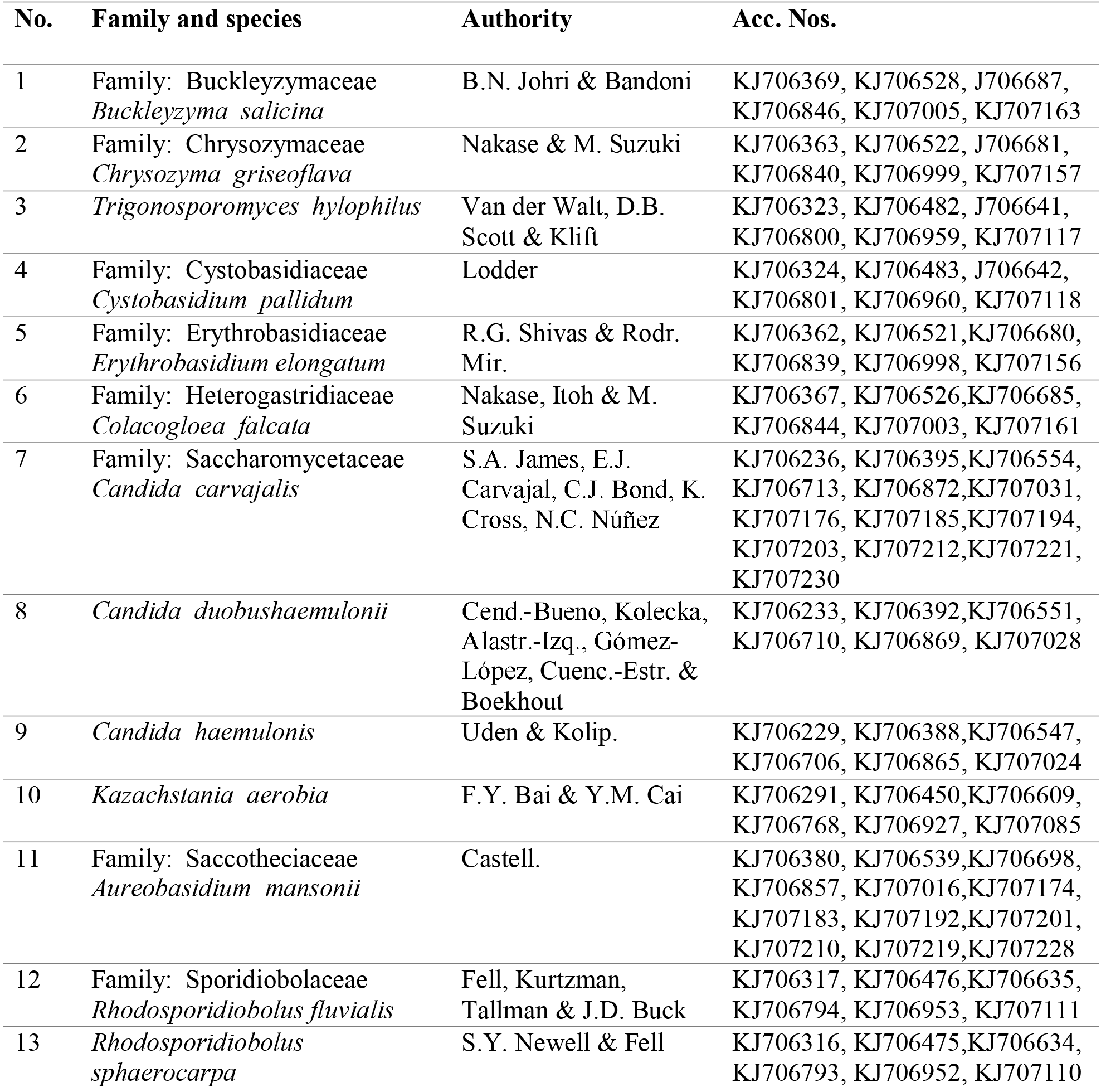
List of marine derived yeast species first time barcoded with their respective species authority and corresponding accession numbers in GenBank

Of the 13 first time barcoded species, a minimum of 0.25 K2P distance and 0.024 nucleotide diversity were reported for *Candida carvajalis* (n = 13) (fig. 2). Maximum K2P distance of 0.37 was recorded for *Buckleyzyma salicina* (n=6), *Trigonosporomyces hylophilus* (n=6) and *Rhodosporidiobolus sphaerocarpa* (n=6) (where maximum of nucleotide diversity of 0.036 was also recorded).

**Fig. 2:**
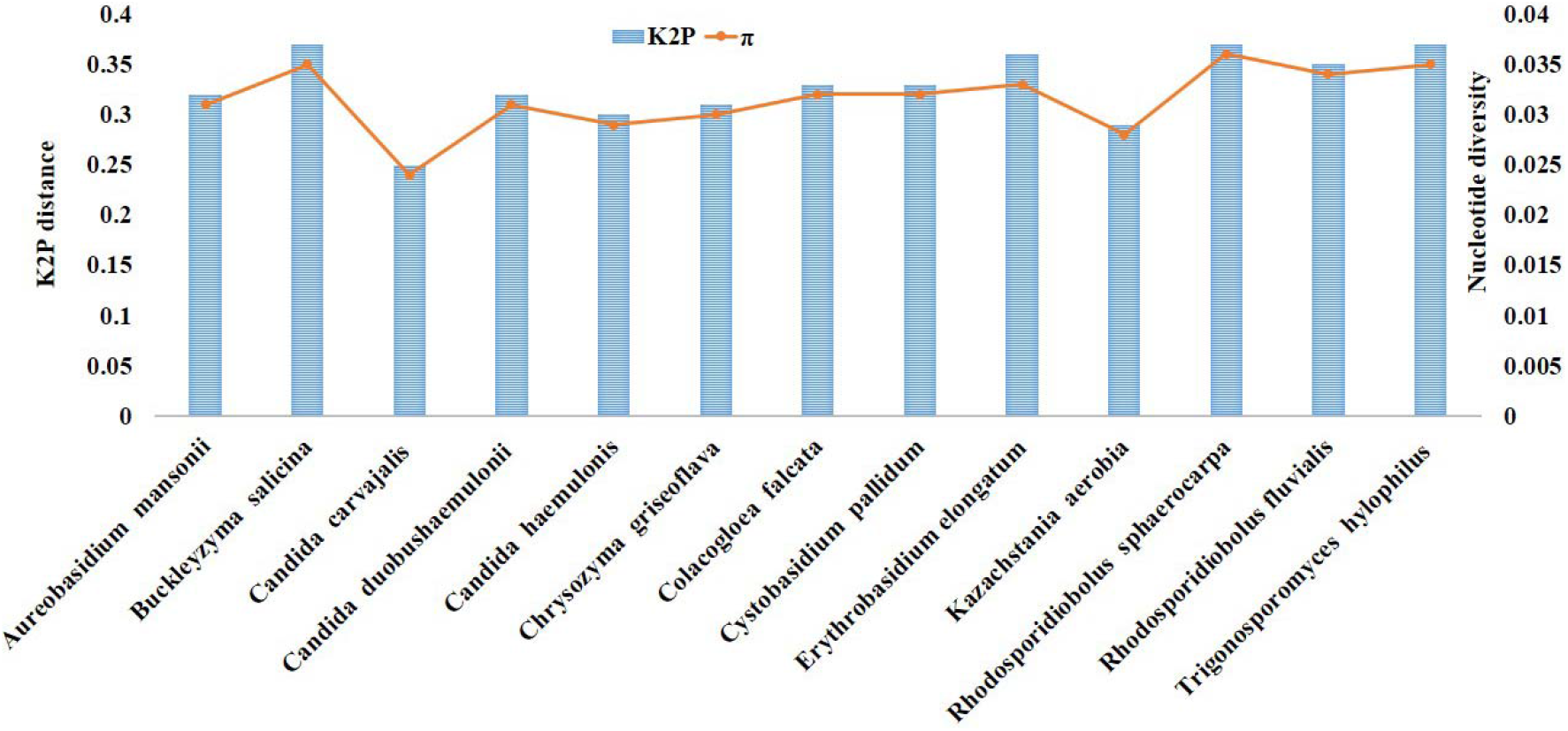
Intra-species average K2P distance and nucleotide diversity (π) of ITS gene sequences of 13 species barcoded for the first time.

The overall intra-species K2P distance average was 0.33% which was well below the threshold proposed for the yeast species (1.59%) (Vu et al., 2016). Nucleotide diversity values directly proportionated the K2P distances (Fig. 2).

### 3.3. Tree based identification

Since the tree based identification was not feasible for all 157 species and requires the construction of mega tress which will be difficult to curate. Randomly selected genera in phylum Ascomycota (*Cryptococcus* spp.) and Basidomycota (*Colacogloea* spp.) were thus used to evaluate the tree based identification. We used a maximum of 3 sequences per species generated in this study belonging to 5 selective species against the corresponding available GenBank reference sequences to create NJ tree of *Cryptococcus* spp. All selected species in the *Cryptococcus* spp. genera, precisely clustered its corresponding reference sequences in one clade (**Fig. 3**). This indicates the efficacy of ITS gene sequences in delineating yeast species. The overall mean K2P pairwise distance of *Cryptococcus* spp. genera was 0.7% which is well within the proposed yeast species cut off (1.59%) (Vu et al., 2016). Tree based *Colacogloea* spp. identification reveals that the individual species clusters the in one clade along with its corresponding GenBank reference sequence (**Fig. 4**). The overall kimura-2 parametric distance was 1.4% which is well within the cut-off value proposed by Vu et al. (2016).

**Fig. 3:**
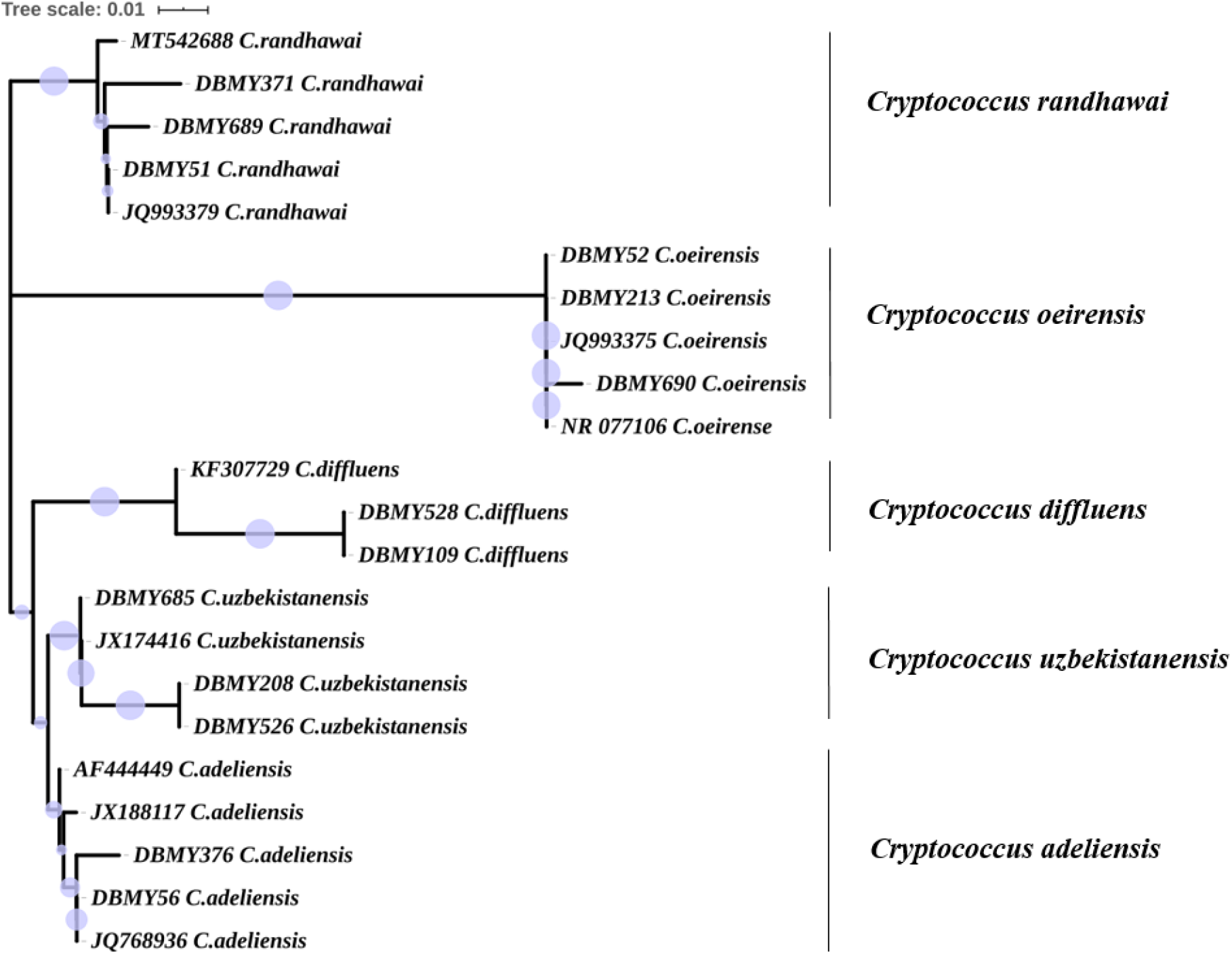
NJ tree based identification of *Cryptococcus* spp. spp. (phylum: Ascomycota, class: Saccharomycetes). The acronym “DBMY” denotes the sequences produced in the present study. The acronym other “DBMY” denotes the accession number of GenBank reference sequences.

**Fig. 4:**
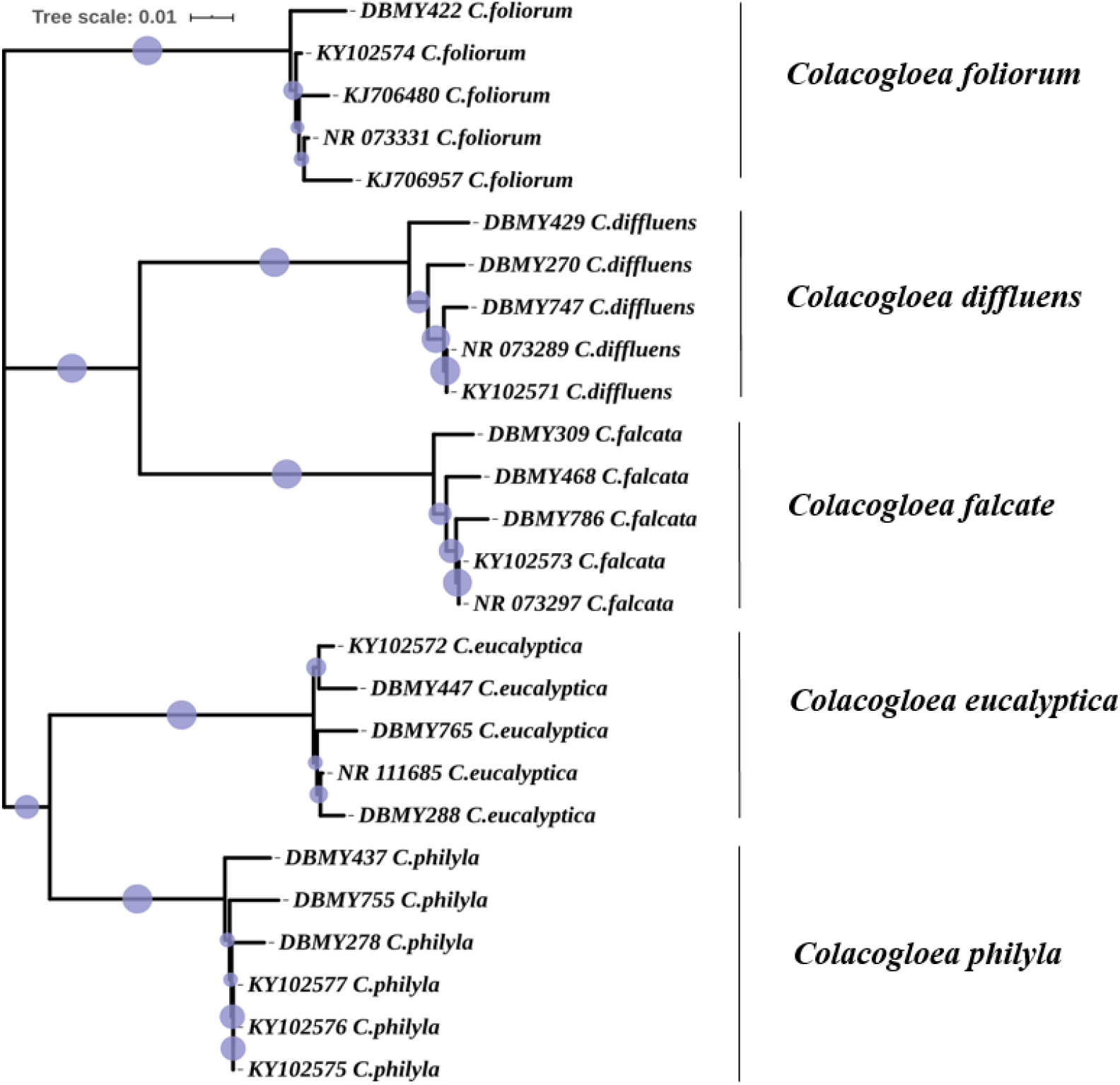
NJ tree based identification of *Colacogloea* spp. (phylum: Basidiomycota, class: Microbotryomycetes). The acronym “DMBY” denotes the sequences produced in the present study.

### 3.4. New occurrences of marine derived yeast species

Only 25.5% (n=40) of the isolated marine yeast species in this study were previously isolated form marine environment. Example; *Sterigmatomyces halophilus* was known to improve marine fish immunity (Reyes-Becerril et al., 2017). *Yarrowia lipolytica* has been known for its crude oil degradation capability (Hassanshahian et al., 2012) and for its dimorphic growth when it is especially isolated from oil polluted seawater (Zinjarde et al., 1998).

Another species isolated in this study, *Candida oceani*, first isolated from hydrothermal vents in Atlantic (Burgaud et al., 2011), was noted for its ability to withstand high hydro-static pressure (Burgaud et al., 2015). *Bandonia marina* reclassified from *Candida marina* (Liu et al., 2015), first isolated from the marine environment (Van Uden and Zobell, 1962), was also recognized for its hydrocarbonoclastic potential (Itah and Essien, 2005), and was previously isolated from tar balls obtained from the northwest coastal waters of India (Shinde et al., 2017). Other species (~58%; n=91) recorded in this study were either reported to occur in soil, plants and animals (including insects) and their presence in the marine environment was previously unrecorded. Approximately 16.6% of the yeast species isolated in this study were previously reported as potential human pathogens (n=26 species).

#### 3.4.1. Yeast species previously unrecorded from marine environment

Approximately 60% of the yeast species (n=94) reported in this study were previously unrecorded in the marine environment. Since it would be exhaustive to describe the previous source of occurrences of 94 yeast species, we present a few examples as follows. *Cystobasidiopsis lactophilus* has been reclassified from *Sporobolomyces lactophilus* (Wang et al., 2015), previously isolated from phyllosphere of the coniferous trees (Nakase et al., 1990). Previously known to occur in forest soils (Mašínová et al., 2018), *Oberwinklerozyma yarrowii* reclassified from *Rhodotorula silvestris* (Wang et al., 2015) was first recorded to occur in mangrove sediments. River run off could be an important medium of transport to occur in mangrove sediment. *Slooffia cresolica* reclassified from *Rhodotorula cresolica* (Wang et al., 2015) was considered to be a part of soil microbiome (Middelhoven and Spaaij, 1997) and its abundances was also correlated with soils with high oil contaminations (Csutak et al., 2005). Formerly known for high hydrocarbon levels (Lyla et al., 2012) are the continental shelf sediments from which these strains were isolated in the present study. Similarly, in the present study *Candida catenulata* previously isolated from polluted sites with hydrocarbon (Habibi et al., 2017; Babaei et al., 2018) was also isolated.

First isolated from tree associated beetles (van der Walt et al., 1971), is *Trigonosporomyces hylophilus* reclassified from *Candida hylophila* (Wang et al., 2015) and their isolation from the mangrove sediments in this study suggests their potential occurrences in mangrove habitat related insects. *Udeniozyma ferulica* and *Vonarxula javanica*, reclassified from *Rhodotorula ferulica* and *Rhodotorula ferulica*, respectively (Wang et al., 2015) were known to occur in polluted river waters (Sampaio and Van Uden, 1991). River run offs could be the reason for these species to occur in mangrove and continental shelf sediments.

The fact that *Debaryomyces mycophilus* was first isolated from wood lice (Thanh et al., 2002) opens the possibility that this species may also occur in mangrove habitat related insects, as the genetic materials of insect could be obtained and studied from the sediment of its habitat (Thomsen et al., 2009). *Debaryomyces pseudopolymorphus* has been extensively involved in the wine fermentation and associated processes (Potgieter, 2004; Villena et al., 2006; Arevalo-Villena et al., 2007). *D. pseudopolymorphus* isolation and its function in the mangrove environment is new and unknown. *Scheffersomyces shehatae* has been known to occur in degrading woods (Kordowska-Wiater et al., 2017) and in wood digesting insects (Suh et al., 2013) was also commonly used for the production of bio-ethanol (Tanimura et al., 2015; Kordowska-Wiater et al., 2017). The association of *S. shehatae* with mangroves and its associated insects could be further explored. *Colacogloea falcatus* reclassified from *Sporobolomyces falcatus* (Wang et al., 2015) were first isolated from dead plant leaves (Nakase et al., 1987). Also, they were isolated from plants phylosphere (Nakase et al., 2003; Takashima and Nakase, 2000) and acidic soils (Delavat et al., 2013). Though ubiquitous distribution trends of certain yeast taxa in decomposing leaves were known (Sampaio et al., 2007), their presence in sediments of mangrove and continental shelf was unknown until now.

*Pichia guilliermondii* has widespread occurrences such as plant endophytes (Zhao et al., 2010), citrus fruit flora (Arras et al., 1998), beetle associated (Suh et al., 1998), and in sewage sludge (de Silóniz et al., 2002). Therefore their isolation in this study may be correlated with multiple sources. *Kazachstania aerobia*, first isolated from plants (Magalhaes et al., 2011) and latter recognised as plant associated yeast (Lu et al., 2004) was unknown to occur in mangrove related habitats until now. *Sporobolomyces koalae* was first isolated from koalas bear (Satoh and Makimura, 2008), and found in other animals such as horses (Fomina et al., 2016) were unknown to occur in marine related habitats until now.

#### 3.4.2. Potential human pathogenic yeasts

Of 157 species isolated in this study, 26 yeast species were recognised as potential human pathogens (**Table S3**). *Candida* spp. contributes 42.3% (n=11 species), followed by *Cryptococcus* spp. (19.2%; n=5). While few human pathogens (such as *Candida orthopsilosis, C. viswanathii, Rhodotorula mucilaginosa* and *Sterigmatomyces elviae*) isolated from this study was previously reported in marine environments (Li et al., 2010; Zaky et al., 2016; Pinheiro et al., 2018; Rasmey et al., 2020), species such as *Candida dubliniensis, C. duobushaemulonii, C. haemulonii, C. nivariensis, C. parapsilosis, C. rugosa, Debaryomyces fabryi* and *Yamadazyma triangularis* were obligate human pathogens (Gasparoto et al., 2009; Ramos et al., 2018; Gade et al., 2020; Ben-Ami et al., 2017; KLi et al., 2014; Mesini et al., 2020; Mloka et al., 2020; Tafer et al., 2016; Kurtzman et al., 2011) with previously unknown occurrences outside of the human host.

## 4. Conclusion

Although the definition of species was not widely applicable (Wheeler and Meier, 2000), in particular for non-obligatory sexually reproductive species such as fungi, the identification of species is a key step for various biological fields such as ecology, agriculture, biotechnology and medicine to identify biological interactions, for example; biodiversity assessment, bioremediation and pathology (de Queiroz, 2007). There could be ~3.8 million unknown fungal species (Hawksworth and Lücking 2017) and the environmental selection pressure plays a crucial role in new species evolution (Handelsman, 2004, Hibbett, 2016; Gabaldón, 2020). The present study was first of its kind in exploring large scale of marine environments for culturable species of yeast. As a result, 1017 barcodes of 157 marine yeast species were produced, of which 91 barcodes of 13 species was barcoded for the first time. This study recorded terrestrial yeast species introduced into the marine environment (ex., *Cystobasidiopsis lactophilus, Oberwinklerozyma yarrowii*) and marine endemic species whose occurrences was restricted to specific marine ecosystem (Ex., *Bandonia marina, Candida oceani*). The DNA barcodes have been published via GenBank and BOLD databases for public use, which will also improve the yeast species barcode coverage and taxonomy in the public databases. The barcodes published in this study (on 2014, in GenBank), have already proved worthy in identifying majority yeast species in previous studies (Vu et al., 2016). These DNA barcodes can also help identify and estimate marine yeast diversity from environmental samples, as many metagenomic diversity studies suffers from lack of local species barcode library (Hawksworth, 2001; Handelsman, 2004; Hibbett, 2016). The entire DNA barcode set produced from this study could be exclusively accessed at http://dx.doi.org/10.5883/DS-MYIC.

Our next challenge will be to explore the synthetic biology, biochemistry, clinical and industrial potential of the isolated strains by venturing into new marine environments with continuous expansion of the barcode databases, as marine derived yeast species were known for many unique features (Guaragnella et al., 2013; Zaky et al., 2014; Dai et al., 2014; Deparis et al., 2017; Zhang et al., 2017; Yashiroda & Yoshida, 2019; Kumar & Kumar, 2019). This may be the largest DNA barcode dataset for culturable marine yeast species. The yeast barcode data produced may be used to explore taxonomic distribution of specific physiological traits (ex., theromotolerance), species of climate and pathological significance (Robert et al. 2015). Correlation of the yeast barcode data with other traits such as the ability to produce various metabolites and industrial products of biotechnological significance (example, antibiotics) would be a valuable resource for yeast researchers willing to apply DNA barcoding technology beyond taxonomic and identification applications.

## Acknowledgement

First author thanks the Department of Science and Technology’s INSPIRE Fellowship (IF10431), India and China Postdoctoral Research Foundation’s National Postdoctoral fellowship (0050-K83008), China for the financial assistance. The authors are thankful for the ship time (FORV-Sagar Sampada) provided by the Centre for Marine Living Resources and Ecology, Ministry of Earth Sciences, Kochi, Government of India for sampling continental shelf sediments under Benthic Productivity program. The encouragement and support provided by Lyla P S and Ajmal Khan S is acknowledged. The authors gratefully acknowledges the Faculty of Pharmacy, and Chiang Mai University, Chiang Mai, Thailand. The research was partially supported by Chiang Mai University.

