## Supplementary table 3 for "Previously unrecorded distribution of marine sediments derived yeast isolates revealed by DNA barcoding"

**Table S3:** List of previously identified, human pathogenic yeast species isolated

| S. No | Species | Clinical sample | Environmental sample |
| --- | --- | --- | --- |
| 1 | *Cryptococcus adeliensis* | Isolated from clinical samples (Tintelnot and Losert, 2005) and causes meningitis (Rimek et al., 2004) | Antartic algae (Scorzetti et al., 2000) and frequently from antartic terrestrial and marine samples (Duarte et al., 2013) including Antartic moss (Zhang et al., 2014) |
| 2 | *Cryptococcus albidus* | Isolated from immune compromised patients (Sugita et al., 2001) and other clinical samples (Lin et al., 1989; Lee et al., 2004) | lake waters (Vadkertiova and Slavikova, 1995) and food samples (Senses-Ergul et al., 2006) |
| 3 | *Cryptococcus diffluens* | Causes subcutaneous cryptococcosis (Serda Kantarcioğlu et al., 2007) and dermatitis in humans (Sugita et al., 2003) | Petroleum contaminated environments (Yilmaz and Sayar, 2015; Yalçın et al., 2014) |
| 4 | *Cryptococcus magnus* | isolated from leukemia patients (Khan et al., 2011) and in other human clinical samples (Ghajari et al., 2018) | Isolated from grapes (Pantelides et al., 2015), plant leaves (Han et al., 2017) and from some cereals (Han et al., 2015) |
| 5 | *Cryptococcus uzbekistanensis* | isolated from immune compromised patients (Powel et al., 2012) | Plant flowers (Hyun and Lee, 2014), and known causative agents of canker in fruit trees (Dehghan-Niri et al., 2015). Also isolated from petroleum contaminated sites (Yalçın et al., 2014; Yılmaz et al., 2015) |
| 6 | *Naganishia globosa* | Clinical samples (Pakshir et al., 2018) | Soil (Kim et al., 2018), hyper saline soils (Mokhtarnejad et al., 2016) and lakes (Li et al., 2018) |
| 7 | *Naganishia liquefaciens* | Isolated from human skin (Timm et al., 2020) and from hospital environments (de Oliveira Brito et al., 2019) | Soil (Abu-Mejdad et al., 2019), lakes (Li et al., 2018) and from municipal wastes (Selvakumar and Sivashanmugam, 2018) |
| 8 | *Pichia farinosa* | Isolated from clinical samples (Adler et al., 2007; Magalhaes et al., 2015) | Halotolerence (Suzuki and Nikkuni, 1989) |
| 9 | *Candida albicans* | Numerous clinical samples (Matthews et al., 1987; Asakura et al., 19991; Nolte et al., 1997; Baena-Monroy et al., 2005;  Dunkel et al., 2008) | Textile effluents (Vitor and Corso, 2008) |
| 10 | *Candida dubliniensis* | Fungal pathogen (Gutierrez et al., 2002), isolated from numerous clinical samples (Chavasco et al., 2006; Barros et al., 2008; Marcos-Arias et al., 2009; Gasparoto et al., 2009) | - |
| 11 | *Candida duobushaemulonii* | Emerging pathogen (Boatto et al., 2016; Frías-De-León et al., 2019) and was recently isolated from numerous cases of clinical samples (Fang et al., 2016; Ramos et al., 2018; Gade et al., 2020) | - |
| 12 | *Candida haemulonii* | Fungal pathogen (Almeida et al., 2012) and was commonly isolated from clinical samples (Rodero et al., 2002 Khan et al., 2007; Ben-Ami et al., 2017) | - |
| 13 | *Candida nivariensis* | pathogen (Alcoba-Flórez et al., 2005; Borman et al., 2008) frequently isolated from various clinical samples (Wahyuningsih et al., 2008; Lockhart et al., 2009; KLi et al., 2014) | - |
| 14 | *Candida palmioleophila* | Fungal pathogen (Sugita et al., 1999; Jensen et al., 2011; Pierantoni et al., 2020) | Environmental samples (Nakase et al., 1988), able to decolorize azo dyes (Jafari et al., 2013) |
| 15 | *Candida parapsilosis* | Fungal pathaogen (Levy et al., 1998; Tay et al., 2009; Canton et al., 2011; Mesini et al., 2020) | - |
| 16 | *Candida rugosa* | fungal pathogen (Pfaller et al., 2006), frequently isolated from various clinical samples (Chaves et al., 2013; Seyom et al., 2020; Mloka et al., 2020) | - |
| 17 | *Candida orthopsilosis* | Numerous clinical samples (Yong et al., 2008; Asadzadeh et al., 2009; Canton et al., 2011;  Feng et al., 2012) | Marine environment (Rasmey et al., 2020) |
| 18 | *Candida spencermartinsiae* | Human pathogen (Shokohi et al., 2018) | Apple fruits (de Garcia et al., 2010) and mangrove ecosystems (Statzell-Tallman et al., 2010) |
| 19 | *Candida viswanathii* | Fungal pathogen (Viswanathan and Randhawa, 1959), Various clinical samples (sandhu et al., 1976; Zaragoza et al., 2011; Wong et al., 2020) | Marine samples (Zaky et al., 2016) |
| 20 | *Aureobasidium mansoni* | Clinical samples (Krcmery et al., 1994) and were recognised as an emerging pathogens (Bamadhaj et al., 2015) | - |
| 21 | *Rhodotorula dairenensis* | Clinical samples (Cobo et al., 2020) | Flowers and fruits of tropical forest (Colla et al., 2010) |
| 22 | *Rhodotorula mucilaginosa* | Clinical samples (Galán-Sánchez et al., 1999; 2011; Da Cunha et al., 2009; Falces‐Romero et al., 2018) | Marine environment (Li et al., 2010) |
| 23 | *Sterigmatomyces elviae* | Human skin lesions (Sonck, 1969) | Guts of fishes (Pinheiro et al., 2018) |
| 24 | *Filobasidium uniguttulatum* | Meningitis in humans (Pan et al., 2012) | Isolated from decaying woods (Jimenez et al., 1991) |
| 25 | *Debaryomyces fabryi* | Human mycotic lesions (Tafer et al., 2016) | - |
| 26 | *Yamadazyma triangularis* | Isolated from human lung tissue (Smith & Batenburg-Van der Vegte, 1986; Kurtzman et al., 2011) | - |
